# Bayesian estimation of MSM population size in Côte d’Ivoire

**DOI:** 10.1101/213926

**Authors:** Abhirup Datta, Wenyi Lin, Amrita Rao, Daouda Diouf, Abo Kouame, Jessie K Edwards, Le Bao, Thomas A Louis, Stefan Baral

**Affiliations:** Department of Biostatistics, Johns Hopkins University; Division of Biostatistics and Bioinformatics, University of California, San Diego; Department of Epidemiology, Johns Hopkins University; Enda-Sante, Dakar, Senegal; Ministry of Health, Cote d’Ivoire; Department of Epidemiology, University of North Carolina, Chapel Hill; Department of Statistics, Penn State University

**Keywords:** AIDS, Bayesian model, Cˆote d’Ivoire, Gaussian process, HIV, MSM population, spatial model, West Africa

## Abstract

Côte d’Ivoire has one of the largest HIV epidemics in West Africa with around half million people living with HIV. Key populations like gay men and other men who have sex with men (MSM) are often disproportionately burdened with HIV due to specific acquisition and transmission risks. Quantifying the MSM population sizes at subnational level is critical to improving the HIV prevention interventions. While survey-based direct estimates of MSM numbers are available at a few urban centers in Cˆote d’Ivoire, no data on MSM population size exists at other areas without any community infrastructure to facilitate sufficient access to the MSM community. We use this limited data in a Bayesian regression setup to produce first empirically calculated estimates of the numbers of MSM in all areas of Cˆote d’Ivoire prioritized in the HIV response. Our hierarchical model imputes missing covariates using geospatial information and allows for proper uncertainty quantification leading to meaningful confidence bounds for the predicted MSM population size estimates. The intended impact of this process is to increase uptake and use of high quality, comprehensive epidemiologic and interventional data in program planning. These estimates will help design future surveys and support the planning of the scale and content of HIV prevention and treatment programs for MSM in Cˆote d’Ivoire.

## 1 Introduction

The last five years have witnessed significant advancements in the response to HIV including universal treatment for those living with HIV, antiretroviral based pre-exposure prophylaxis to prevent HIV, and new diagnostic approaches including HIV-self testing (UNAIDS 2017). However, leveraging these strategies to achieve an AIDS-Free generation by 2030 necessitates understanding who and why people continue to acquire HIV (Beyrer et al. 2014, Stahlman et al. 2016). In concentrated epidemics, it has long been understood that the majority of HIV infections are among populations with specific acquisition and transmission risks for HIV including gay men and other men who have sex with men (MSM), sex workers, people who use injection drugs, and transgender women (Beyrer et al. 2014). However, in generalized HIV epidemics, the specific proximal risks for HIV have been less studied which challenges the ability to effectively specify both the most appropriate bene-factors for these new interventions as well as the number of people in need (Boily et al. 2015, Mishra et al. 2016). To address the former, there has been a number of epidemiologic and mathematical modeling studies demonstrating the importance of addressing the HIV prevention and treatment needs of key populations in the context of generalized HIV epidemics (Mishra et al. 2016). However, there remain limited data on the sizes of key populations across most generalized HIV epidemic settings (Abdul-Quader et al. 2014).

Characterizing the numbers of key populations facilitates an understanding of the numbers of potential eligible candidates for more intensive HIV prevention interventions, the overall potential impact of those interventions when implemented at scale, and finally an improved understanding of the local HIV epidemic (Abdul-Quader et al. 2014, Holland et al. 2016). Moreover, to effectively parameterize mathematical models characterizing the modes of transmission, high quality data regarding the size, characteristics, and HIV burden among key populations are needed (UNAIDS/WHO Working Group on Global HIV/AIDS and STI Surveillance 2010). Concurrently, there has been increasing consensus on the appropriate methods for population size estimation for key populations (Quaye et al. 2015, UNAIDS/WHO Working Group on Global HIV/AIDS and STI Surveillance 2010). However, many current size estimates that have been completed resulted in national estimates, with less in the literature focused on subnational estimates in the majority of low and middle income settings (Tanser et al. 2014). Ultimately, size estimates at the subnational level are those most often used by by local ministries of health, implementing partners, and bilateral and multilateral funding agencies to guide the geographic and population prioritization of resources and efforts (Yu et al. 2014). Often the direct estimates of key population size have been in urban or peri-urban areas where the population densities of key populations are higher and where the community infrastructure exists to facilitate sufficient access to the community (Yu et al. 2014). However, HIV prevention and treatment needs are universal, necessitating methods for estimating population size of key populations at high risk of HIV acquisition and transmission at the subnational level (Tanser et al. 2014).

There exist a range of extrapolation methods to generate estimates at the national and subnational level. These methods differ in terms of their reliance on data, cost, and scientific rigor (Yu et al. 2014). Expert opinion involves consulting experts, including national stakeholders, technical experts, and key population groups, on how confident they are with the direct estimates and seeking their advice on how to apply these numbers to other off-sample areas. This method has low reliance on data, little cost, and relatively low scientific rigor. Simple and stratified imputation apply the mean from areas with direct estimates to the areas where predictions are needed. These methods have some reliance on auxiliary data and result in arguably more evidence-based rigor than relying on expert opinion alone. Less is known about other more complex methods, including regression, treating off-sample areas a missing data problem, and utilizing geospatial covariation or correlation to predict values, i.e., small-area estimation.

In West Africa, the epidemiology of HIV has been distinct from that in Eastern and Southern Africa (Djomand et al. 2014, Holland et al. 2016, Papworth et al. 2013). Specifically, the population HIV prevalence among all adults has not surpassed 5% though very high burdens have been observed among key populations Djomand et al. (2014). The burden of HIV in Cˆote d’Ivoire was estimated to be 3.2% among all adults equating to nearly half a million people living with HIV, nearly all of whom are over fifteen years old. In the national strategic plan for HIV, key populations including MSM, female sex workers (FSW), and people who inject drugs (PWID) have been deemed to be priority populations for HIV prevention and universal treatment for those living with HIV. However, similar to other settings, the enumeration and representative sampling of key populations has been challenged by the criminalized nature of these populations combined with high levels of stigma (Beyrer et al. 2012). Consequently, specialized sampling strategies for key populations in these settings have been used including respondent-driven sampling, time-location sampling, the prioritization for local AIDS control efforts (PLACE) and others. However, the majority of these studies have taken place in urban centers resulting in limited study of population size estimates for key populations in the majority of the country including rural, less densely populated settings Abdul-Quader et al. (2014).

Given limited data on population size of key populations in Cˆote d’Ivoire, the objective of this study was to assess the utility of small area estimation approaches to estimate the population size in the organizational units prioritized by the Presidents Emergency Plan For AIDS Relief (PEFPAR) along with proper quantification of uncertainty of those estimates. Specifically, the study aimed to utilize available direct estimates of population size and other demographic covariates in Cˆote d’Ivoire to generate model based estimates of population sizes of MSM for subnational areas.

## 2 Data

The size of MSM population was directly estimated for five regions of Cˆote d’Ivoire: Abidjan, Agboville, Bouake, Gagnoa, and Yamoussoukro from March 2015-February 2016. Regions for direct size estimation were selected to coincide with a respondent-driven sampling study of adult MSM in progress in these same five areas.

### 2.1 Direct estimates

Size estimates were generated through the use of various multiplier methods, including the unique object multiplier, NGO membership multiplier, service multiplier, and social event multiplier. Briefly, multiplier methods for size estimation compare two independent sources of count population size, assessing the overlap between these independent estimates in order to determine the total number of individuals in the population. The basis for these methods rests on the understanding that the proportion of individuals in the population who appear at a specific institution during a certain time period, for example a current registry of NGO membership, is equal to the proportion who appear at that same institution among the survey participants.

The first independent source of data consisted of a count or listing from programme data for NGO membership, service provision, and social event attendance. For the unique object multiplier method, the first source of data was a log of how many objects were distributed. The total number of MSM attending services at “Clinique de Confiance” was captured from program logs. The total number of MSM who attended the social event “evening GNARA” was recorded. The total number of MSM belonging to NGOs was also captured from program logs.

The second independent source of data for our estimates was a representative survey in which MSM were recruited through Respondent Driven Sampling (Heckathorn 1997, RDS), which is a strategy employed when individuals in the target population are hard-to-reach and when no known sampling frame exists. Methods for respondent-driven sampling have been described previously. Individuals were purposively asked questions to generate this second independent source of data for size estimation (Appendix A). As one example, for the service multiplier method, participants were asked if they had ever received services from “ Clinique de Confiance.”

From these two independent sources of data, direct estimates were generated using the formula *S* = (*N*_1_ *× N*_2_)*/R*, where *S* is the estimate of total population size, *N*_1_ and *N*_2_ respectively are the total numbers of people captured in the first and second independent source of data (e.g. programme log and MSM recruited to the cross-sectional RDS study), *R* is the overlap between two independent sources of data (e.g. number of MSM participating in cross-sectional RDS study who reported accessing services). Since, the estimates were based on RDS, 95% confidence intervals could be calculated (Salganik 2006) for the proportion *R/N*_2_ of RDS participants who were also enlisted in the first programme. Subsequently they are used to create confidence intervals for *S*.

Because those sampled were 90% from the age group 18–29, population size estimates were age-standardized to get a better estimate of all men (15–49) who have sex with other men. In order to age-standardize, a population size estimate proportion was calculated using the male population 18–29 of the given region. In accordance with previous studies looking to estimate the size of men who have sex with men (Purcell et al. 2012), it was assumed that the proportion of MSM remains constant across age groups. Therefore, the calculated population proportion was applied to the total male population 15–49 to get an estimate of the total number of men 15–49 who have sex with other men. However, for the data analysis we used the direct estimates for the 18–29 age group, instead of the age-standardized version for the 15–49 age group. Hence, the data analysis and subsequent predictions of number of MSM in the age group of 18–29 years does not rely on any such assumption.

Table 1 presents the data for the five regions. Observe that not all survey methods were implemented in all areas. In Abidjan, there were two NGOs where we had access to the total count of members. On the other hand, Agboville does not have a service multiplier based estimate and Gagnoa does not have an estimate from NGO membership.

**Table 1:**
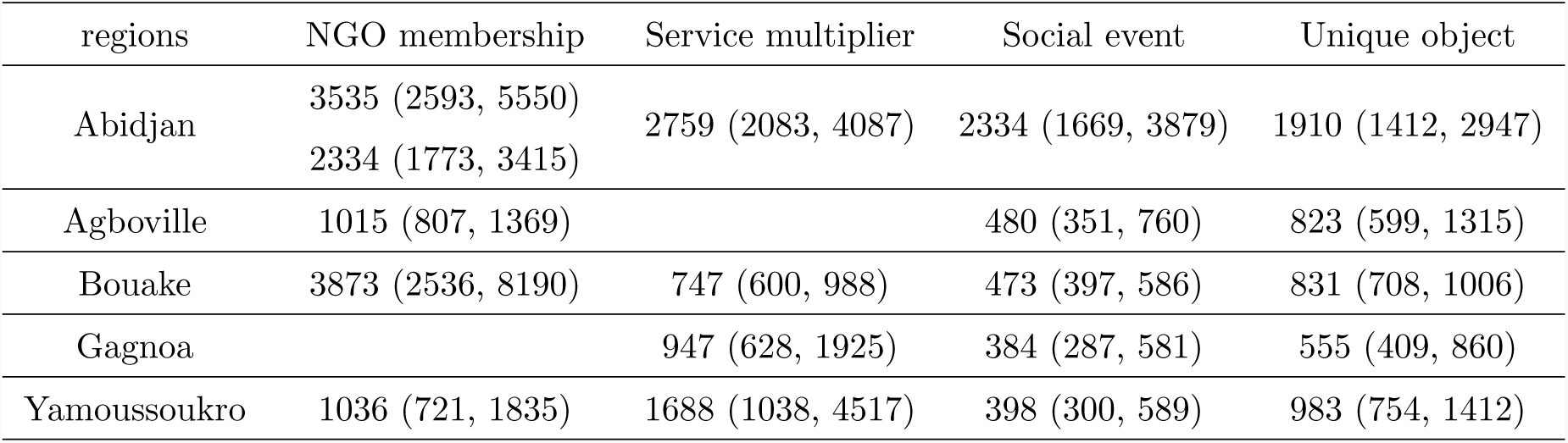
Population size estimates (and 95% confidence intervals) of MSM in age group of 18 to 29 years.

### 2.2 Prediction areas

In total, there were 61 prediction areas that were selected to coincide with PEPFAR’s official Organizational Units (OUs), and that also roughly correspond with Cˆote d’Ivoire’s department-level administrative division. Prediction areas were selected to coincide with PEPFAR Organizational Units (OUs) in order to provide evidence-based estimates for targeted prevention, care and treatment programs and to inform Country Operational Processes. These PEPFAR OUs also roughly correspond with the official department-level administrative division in Cˆote d’Ivoire. In settings where public health systems are decentralized, estimates for program denominators are needed at both the national and sub-national level in order to set actionable targets. This is especially important if there are large regional differences in burden of disease, resources, etc. The intended impact of this process is to increase uptake and use of high quality, comprehensive epidemiologic and interventional data in program planning, while building consensus on small area estimations of available data to guide additional data collection and programmatic efforts focused on HIV among key populations.

### 2.3 Covariates

Covariates were selected based on relevance to prediction model and availability of quality data at the appropriate administrative division (department level). Data for population density, density change and male population was obtained from publicly available data published by the Institut National de la Statistique, Republique de Cˆote d’Ivoire. Data for HIV prevalence was obtained from a UNAIDS report on subnational estimates of HIV prevalence in Cˆote d’Ivoire (http://www.unaids.org/sites/default/files/media_asset/2014_subnationalestimatessurvey_C\unhbox\voidb@x\bgroup\let\unhbox\voidb@x\setbox\@tempboxa\hbox{o\global\mathchardef\accent@spacefactor\spacefactor}\accent94o\egroup\spacefactor\accent@spacefactortedivoire_en.pdf)

There was no department-level age-stratified, sex-stratified data. We assumed a constant age and sex distribution across all departments: 55% of total male population for each of the departments/region seats is in the age group 18–29. Also, for Abidjan, close to 90% of our sample reported being from either Abobo, Cocody, Marcory, Triechville, or Youpougon. This is just five communes out of the total ten in Abidjan. We therefore considered our sample to better represent these five communes of Abidjan rather than the whole city. The total male (15–49) population for these communes was 842551 rather than 1286750 for the whole city and for men 18–29 was 368097 rather than 562160. We also assumed that the age-sex distribution was the same across all the communes.

Additionally, we also used estimated population density from the Landscan database (
http://web.ornl.gov/sci/landscan/) based on night light data. The night light data is from Defense Meteorological Satellite Program (DMSP) Operational Linescan System (OLS) which detects nighttime lights from satellite imagery. Landscan provided estimated population size over 1km*×*1km grid cells. For each of the 61 prediction areas, the population estimates were averaged over a 25 km^2^ radius centered on the area to obtain the average population density for the areas.

## 3 Methods

### 3.1 Linear model

We use the data for the five regions to train a regression model for predicting the MSM population size based on the covariates, and subsequently use this model to extrapolate at all other regions. A lot of our modeling choices are guided by the extremely small sample size (19 datapoints in total from five regions) which proscribed the use of complex models involving many parameters. For the *i*^*th*^ region, let *N*_*i*_ denote the total male population in the age group of 18–29 years, *x*_*i*_ denote the set of demographic covariates and *n*_*ij*_ denote the direct estimate obtained from the *j*^*th*^ method. A natural choice for modeling the population size would have been a generalized linear model (GLM) *n_ij_ ∼* Binomial(*N*_*i*_, *p*_*i*_) where 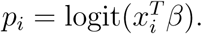 However, note that not all the direct estimates are equally reliable. For example, we observe in Table 1 that the NGO membership based estimate of MSM population in Bouake differs by an order of magnitude from the other three estimates for the same region. The confidence interval for this estimate is also very wide suggesting limited reliability of the estimate. While it is less clear how to incorporate information from the confidence intervals in a binomial GLM setup, we can leverage these data in a linear regression setup via heteroskedastic errors.

Defining *y*_*ij*_ = log(*n*_*ij*_/*N*_*i*_), we specify the linear regression model as 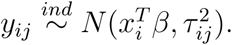 The variance of a normally distributed random variable is proportional to the square of two-tailed 95% coverage interval. Therefore, we specify 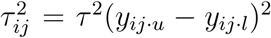 where *y*_*ij*⋅*u*_ and *y*_*ij*⋅*l*_ denotes the upper and lower bounds corresponding to *y*_*ij*_. This ensures that more uncertain estimates with very wide confidence bounds are given less weights in the model than the ones with narrow confidence intervals providing more precise information.

Regression models for small area estimation often include region specific random effects to improve estimation (Fay & Herriot 1979). However, Datta et al. (2011) argued that when the number of regions is small, the simpler model without random effects often performs better. Owing to the very small sample size of the dataset, we decided against introducing region specific random effects as it involves additional parameters.

Finally, we have used a log-transformation to define the *y*_*ij*_’s instead of a logit transformation, although the latter is more natural, as *n*_*ij*_/*N*_*i*_ is a fraction. In the dataset, the proportions *n*_*ij*_/*N*_*i*_ are typically very small (80% are less than 0.05). So the two transformations yield very similar *y*_*ij*_’s. Furthermore, as we discuss in Section 3.2, the log transformation is more interpretable in our final model which includes log(*N*_*i*_) as one of the covariates. Hence, we preferred the log-transformation.

### 3.2 Covariate Selection

The covariates described in Section 2 were region specific. Hence, although there were 19 datapoints, there were only 5 unique sets of covariate values, one for each region. This impeded exploring nonlinear models linking *y*_*ij*_’s to *x*_*i*_ and confined us to the parsimony of the linear model. Even in a linear setup, we want to select only one or two most relevant covariates from the five available — male population, population density, density change, HIV prevalence and Landscan density. We kept HIV prevalence in the model as we expected areas of where there are more MSM to be areas with high HIV prevalence, as MSM are disproportionately affected by HIV given the biology of HIV transmission combined with limited programming focused on mitigating risks specifically among gay men and other MSM. The total male population in the age group (*N*_*i*_) has already been used to define *y*_*ij*_’s. Hence, it seems natural to exclude it from the linear model. However, Figure 1 reveals that *y*_*ij*_’s have a very strong negative correlation with log(*N*_*i*_)’s. Initially this negative correlation seems counter-intuitive given a rural-to-urban migration. One likely explanation is that in large urban centers the MSM community grows at a slower rate than the overall population even if the absolute numbers of MSM is higher. For example, if the numbers of MSM population grows at a rate proportional to 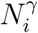 for some *γ <* 1, then cov(*y_ij_,* log(*N*_*i*_)) = *γ −* 1 which is negative. Hence, we included log(*N*_*i*_) as a covariate, which in turn justifies use of a log-transformation to define *y*_*ij*_’s, as it imparts a nice interpretability about of the relative growth rates of the MSM population and the total population. The other explanation would be that in urban areas a higher proportion of MSM are not accounted for in the survey or that the independent assumption is violated in the capture-recapture method. While both reasons are conceivable, the first is a feature of MSM population dynamics while the second is a sampling issue. In the absence of additional data collection, it is not feasible to discern between these two scenarios. However, internet-based surveys may facilitate learning more about the numbers of MSM in more stigmatizing settings.

**Figure 1:**
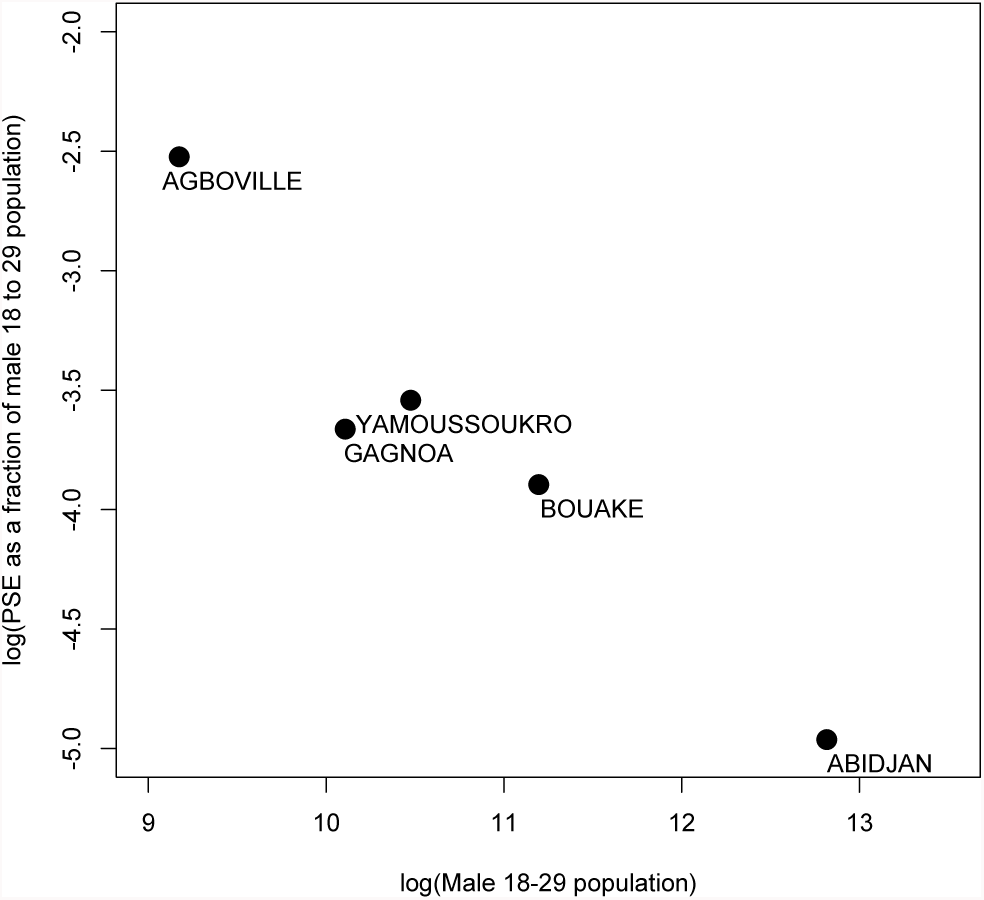
Negative correlation in log-log scale between MSM proportion and total male population of age 18-29.

Due to limited sample size, we restricted model selection only from linear models using no more than two of the five covariates and used leave-one-out cross validation to choose the covariate (alongside HIV prevalence). For each of the 4 models, we leave one region out and fit the model using the remaining regions to predict the MSM population size at the left out region. We repeat this for all the five regions and the average squared error between the predicted estimate and average direct estimate for each region gives the leave-one-out cross validated (LOOCV) error.

Table 2 provides the LOOCV based Mean Square Error for the four models. We found that the model with log male population and HIV prevalence as covariates has the lowest MSE. Thus, the final linear regression model is *y*_*ij*_ = *β*_0_ + *β*_1_ log(*N*_*i*_) + *β*_2_ *H*_*i*_ + *ε*_*ij*_ where *H*_*i*_ denotes the HIV prevalence for the *i*^*th*^ region.

**Table 2:** Leave-one-out cross validated mean square error for the four models

| Covariates included | MSE<sub>LOOCV</sub> |
| --- | --- |
| log(male population) + HIV prevalence | $3.5 \times 10^{-3}$ |
| Landscan density + HIV prevalence | $5.7 \times 10^{-3}$ |
| population density + HIV prevalence | $6.5 \times 10^{-3}$ |
| density change + HIV prevalence | $6.7 \times 10^{-3}$ |

### 3.3 Spatial model for HIV prevalence

HIV prevalence data were missing at around 50 % (30 out of 61) of the locations where we want to predict MSM population. Since, it is one of the covariates in the model, we need to impute the missing values. A simple choice for imputation would be to use the average of the observed values. However, exploratory data analysis using empirical and exponential semivariograms confirms significant spatial pattern (see Figure 2) which can be potentially leveraged to improve the quality of imputation (we refer the reader to the books by Cressie & Wikle 2011, Banerjee et al. 2014, for details on variograms and spatial models).

**Figure 2:**
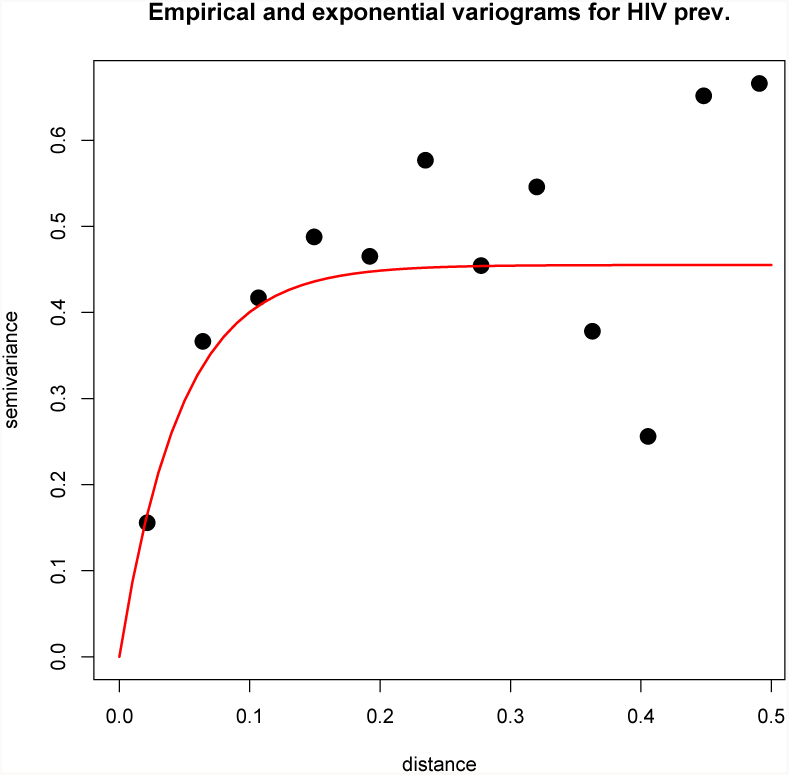
Empirical (black dots) and exponential (red line) variogram for HIV prevalence.

The use of a spatial model was corroborated by a leave-one-out cross validation using the available HIV prevalence data for the 31 regions. Let *s*_*i*_ denote the co-ordinates representing the *i*^*th*^ region and *H*(*s*_*i*_) = *H*_*i*_ denote the corresponding HIV prevalence. We modeled *H*(*s*) as a Gaussian Process (GP) with constant mean and exponential covariance function. If *H*(*S*) denotes the vector formed by stacking up the HIV prevalence data for a set of regions *S*, then the GP specification effectuates a multivariate Gaussian distribution *H*(*s*) *∼N* (*µ*1, Σ) where Σ = *σ*^2^ exp(*−φ || s_i_ − s_j_ ||*)*_s__i,s__j ∈S_*. The prevalence data at *S* is used to estimate the parameters (*µ, σ*^2^*, φ*). Subsequently, the prevalence at any new location *s* is given by

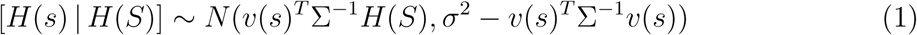

where *v*(*s*) = *σ*^2^(exp *−φ||s − s_i_||*)*_s__i∈S_*. For the cross validation, we use these kriging equations to impute the HIV prevalence at each left out region, based on parameter estimates using data from the remaining regions. For comparison, we used the mean HIV prevalence of the in-sample data to predict at the left out region. The leave-one-out MSE for the spatial model (MSE=0.37) was around 20% better than for the mean imputation (MSE=0.48). Hence, we use the spatial GP model for imputing the missing HIV prevalence data.

### 3.4 Hierarchical Bayesian Modeling

Obtaining meaningful confidence bounds for the predicted MSM population is critical. The uncertainty of the regression parameters and especially the spatial parameters are often ignored in frequentist predictions. Furthermore, for regions with missing HIV prevalence data, the kriging estimates in Equation 1 are accompanied by the kriging variances which can be large if the location is far from the data locations. Hence ignoring this source of uncertainty can lead to narrow prediction bounds. In a frequentist setting, it is unclear how to utilize the kriging variance when the imputed HIV prevalence will be used as a covariate to predict MSM population size. However, we can seamlessly integrate this multistage procedure into a hierarchical Bayesian model which allow for proper propagation of uncertainty associated with all different parts of model estimation into the final prediction of population size estimates.

Let *S* denote the set of locations where HIV data are available. Also, for any location*s*, let *N* (*s*) and *H*(*s*) respectively denote the male population of 18−29 years and the HIV prevalence. Finally, defining *y*_*j*_(*s*_*i*_) = *y*_*ij*_, *w*_*ij*_ = (*y_ij,u_ − y_ij,l_*)^2^ and *β* = (*β*_0_*, β*_1_*, β*_2_)^′^, the full specification of the hierarchical model is given by:

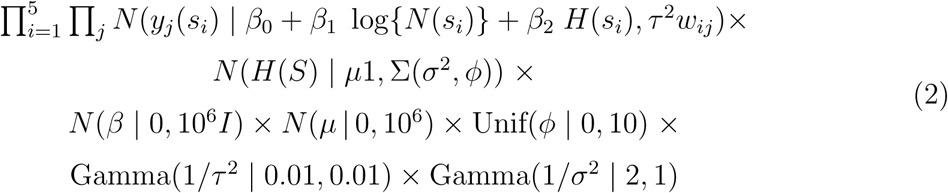

The top row of 2 is the log-normal regression model for the MSM percentages, the middle row is the spatial Gaussian Process model for HIV imputation and the bottom two rows are the parameter priors. Gamma(*a, b*) denotes the Gamma distribution with shape parameter *a* and rate parameter *b* and Unif(*a, b*) is the uniform distribution on (*a, b*). We use the *Nimble* package in R (https:\\r-nimble.org) to generate 30, 000 MCMC samples from this model, the first 15, 000 of which is discarded as burn-in. The posterior estimates for all the parameters are provided in Table 3. We observe that there is strong negative correlation between *y*_*ij*_ and log(*N*_*i*_) which we have discussed in Section 3.2. The association with HIV prevalence is relatively weak. The estimates of the spatial parameters indicate a strong spatial dependence in HIV prevalence, previously insinuated by the variograms in Figure 2.

**Table 3:** Posterior median and 95% credible interval for the hierarchical model

|  |  |  |  |
| --- | --- | --- | --- |
| $\beta_0$ | 2.62 (0.04, 5.23) | $\mu$ | 2.55 (1.54, 3.61) |
| $\beta_1$ | -0.79 (-1.09, -0.51) | $\sigma^2$ | 0.86 (0.5, 2.29) |
| $\beta_2$ | 0.63 (-0.07, 1.33) | $\phi$ | 7.68 (2.71, 9.89) |
| $\tau^2$ | 0.65 (0.34, 1.43) | | |

### 3.5 Prediction

We use composition sampling to obtain posterior predictive distributions of MSM population size at a new location. If *s ∈ S*, then this is simply given by the samples 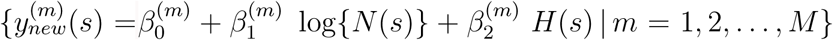 where 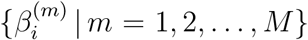 denote the MCMC samples from posterior distribution of *β_i_*. For locations outside *S* with no HIV prevalence data, posterior distribution of *H*(*s*) is given by

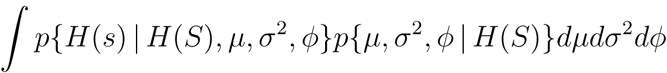

This is effectively accomplished using the samples *{µ*^(*m*)^, (*σ*^2^)^(*m*)^*, φ*^(*m*)^*}* to generate *H*(*s*) *| H*(*S*) via the kriging Equation in (1). Subsequently, the samples 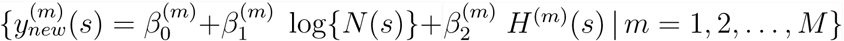 represent posterior predictive distribution for MSM population size at those locations.

We train the linear regression model on a very limited set of values of the predictors, based on just five unique datapoints. It is difficult to assess a priori whether we can extrapolate this linear relationship to other regions with significantly different demographics. We observed that for some areas with very low population, the predicted MSM population percentage was abnormally high. Further investigation into this reveals that the minimum total male population among the five regions with survey data corresponds to the 36^*th*^percentile of the empirical distribution of total male population among all the 61 regions. Hence, the training data corresponds to larger areas with greater population and does not inform much about the regression relationship in regions where the population is very low. This, combined with the strong negative value of *β*_1_ in Table 3 results in such high estimated MSM fractions.

As a heuristic remedy, we assume that the negative relationship flattens out below a certain population threshold. We truncate the total male population at the 10% quantile of the empirical distribution and use these thresholded values for prediction. While this is ad hoc, more formal methods like estimating the truncation point based on the data will always truncate within the data values, whereas replacing a linear regression with a general monotonic function will involve more parameters and hence is infeasible for our small dataset. Of course, our truncation does not affect parameter estimation as all the total population values for the training data are above the truncation level. This issue is less severe for HIV prevalence as the observed values for the five regions better represent the empirical distribution of HIV prevalence. Since, it also has a much weaker association with population size of MSM, we do not truncate the HIV prevalence values.

## 4 Size estimates

Figures 3, 4 and 5 present the uncertainty quantified predictions of HIV prevalence, MSM population fraction and size respectively while the actual numbers are presented in Table 4. For the regions with no direct estimates, the predicted MSM population percentage typically varied between as low as 0.5% to around 10%. The highest MSM percentages are predicted in Katiola, Kouassi-kouassikro and Bettie. However, these areas also had the widest credible intervals indicating the large uncertainties associated with the predictions. Figure 6 demonstrates the impact of HIV imputation on the prediction uncertainties. Since, the variance and width of confidence intervals of log-normal distribution are proportional to the mean, we use relative width (ratio of the 95% cofidence interval width to the estimate) as a more meaningful measure of uncertainty. In Figure 6a we plot the predictions of MSM population percentage against the relative width. We observe that the relative width was in general larger for locations with missing HIV prevalence data. This is expected as the Bayesian model properly propagates the uncertainty associated with the imputation of HIV prevalence in the final predictions. This is nicely reflected in the CI widths. In Figure 6b we plot the relative width against leverages for each region. For regions with HIV prevalence data, the relative widths increase with the leverage as expected indicating that predictions for regions with covariates values distant from those of any of the observed regions are accompanied with larger uncertainty. Among regions with missing HIV prevalence, this trend was less prominent due to the added component in the uncertainty from the imputation.

**Figure 3:**
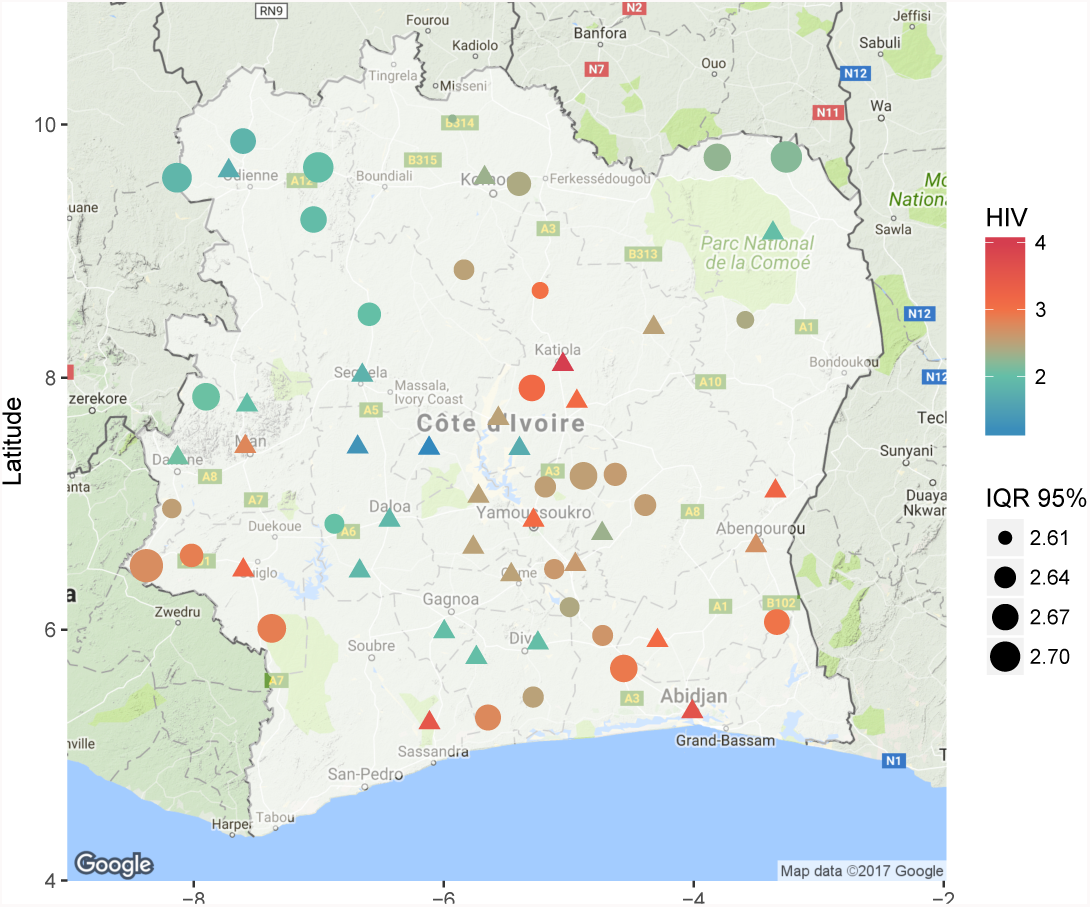
Predicted and observed HIV. ∆ represents observed data. IQR 95% is the 95% Inter-quantile range, i.e. width of the 95% credible interval.

**Figure 4:**
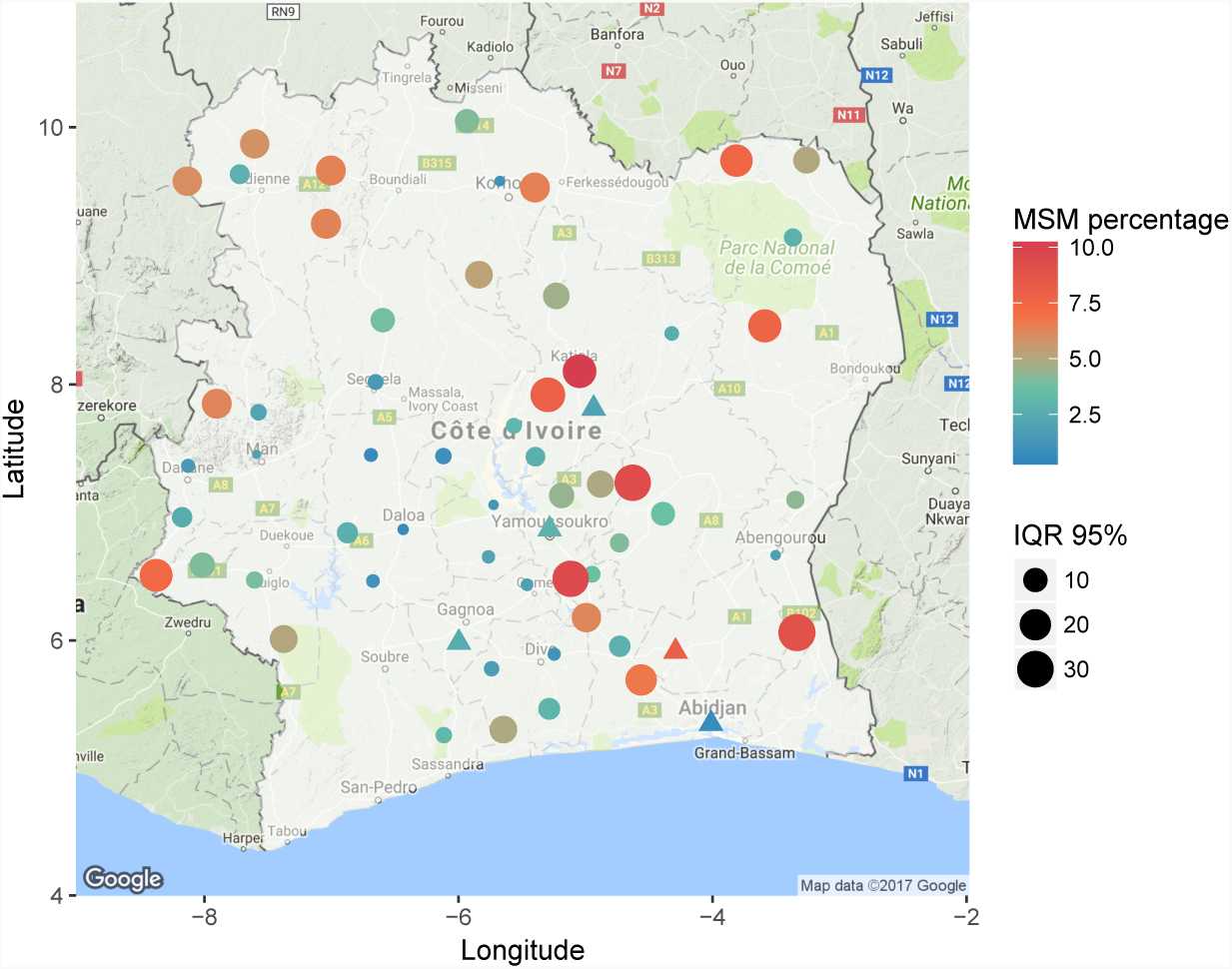
Predicted and observed MSM population as a percentage of total population. ∆ represents observed data. IQR 95% is the 95% Inter-quantile range, i.e. width of the 95% credible interval.

**Figure 5:**
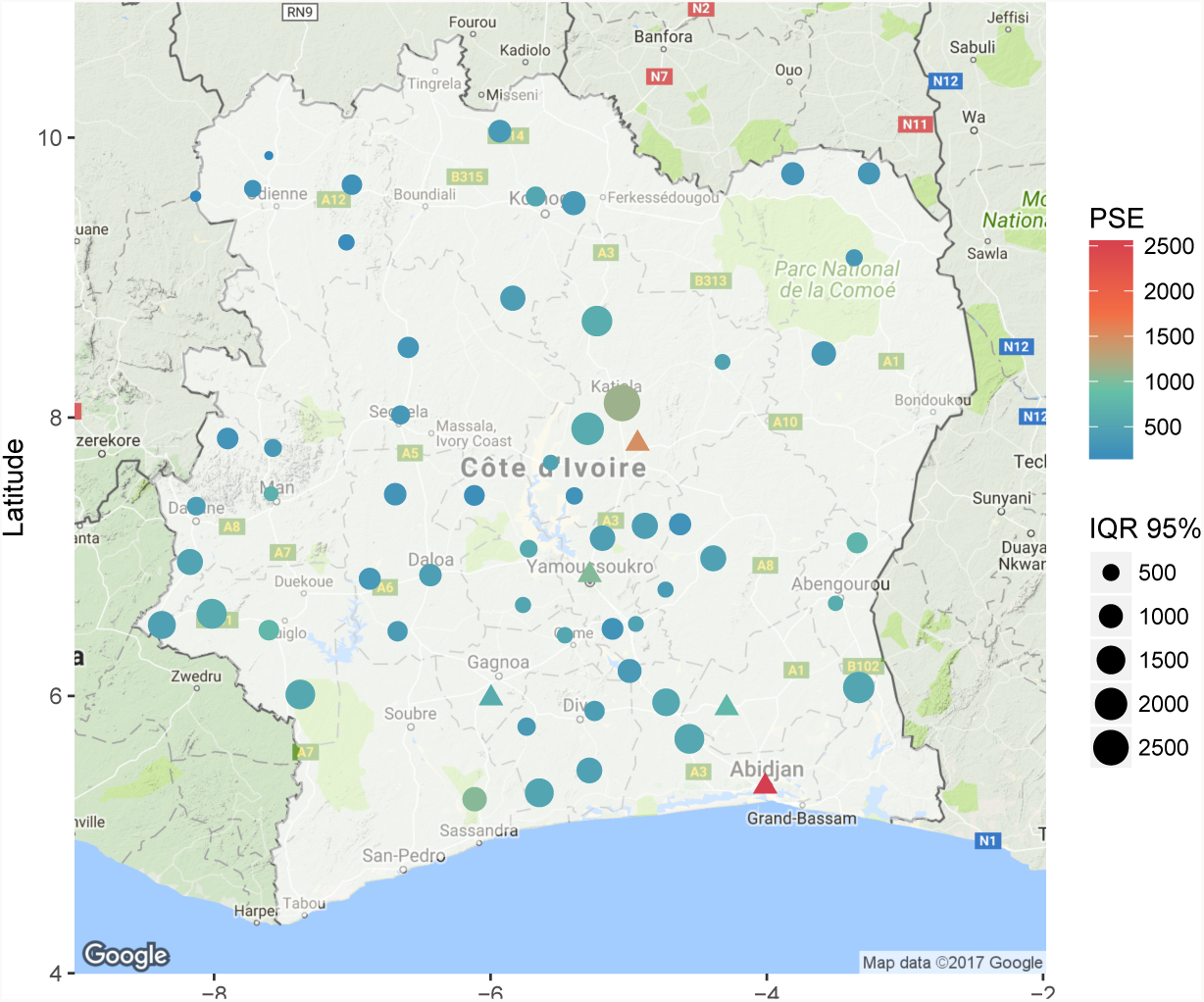
Predicted and observed MSM population. ∆ represents observed data. IQR 95% is the 95% Inter-quantile range, i.e. width of the 95% credible interval.

**Table 4:** Predicted MSM population size, population fraction and HIV prevalence along with credible intervals (within braces). Bold font indicates HIV prevalence for regions were they were observed (these are direct estimates and hence don’t have credible intervals).

| Regions | MSM population size | MSM population % | HIV prevalence |
| --- | --- | --- | --- |
| Abidjan | 1818 (1113, 2941) | 0.5 (0.3, 0.8) | **3.6** |
| Agboville | 628 (352, 1134) | 6.5 (3.7, 11.8) | **3.1** |
| Bouake | 953 (741, 1218) | 1.3 (1, 1.7) | **3.1** |
| Gagnoa | 383 (187, 783) | 1.6 (0.8, 3.2) | **2** |
| Yamoussoukro | 822 (614, 1101) | 2.3 (1.7, 3.1) | **3.1** |
| Abengourou | 638 (457, 889) | 1.8 (1.3, 2.6) | **2.7** |
| Agnibilekrou | 755 (472, 1218) | 4.3 (2.7, 7) | **3.2** |
| Beoumi | 476 (305, 746) | 3.1 (2, 4.8) | **2.5** |
| Biankouma | 356 (176, 721) | 2.1 (1, 4.2) | **2** |
| Bouafle | 591 (377, 922) | 1.3 (0.9, 2.1) | **2.5** |
| Bouna | 328 (164, 666) | 2.8 (1.4, 5.8) | **2** |
| Dabakala | 497 (323, 769) | 2.6 (1.7, 4) | **2.5** |
| Daloa | 439 (182, 1051) | 0.7 (0.3, 1.6) | **1.9** |
| Danane | 421 (216, 816) | 1.5 (0.8, 2.9) | **2.1** |
| Dimbokro | 377 (216, 673) | 4.1 (2.4, 7.3) | **2.3** |
| Divo | 422 (198, 894) | 1.1 (0.5, 2.3) | **2** |
| Guiglo | 772 (495, 1214) | 4 (2.6, 6.3) | **3.2** |
| Issia | 387 (171, 865) | 1.1 (0.5, 2.5) | **1.9** |
| Katiola | 1134 (413, 3078) | 10.3 (3.7, 27.8) | **4** |
| Korhogo | 543 (301, 975) | 1 (0.6, 1.8) | **2.3** |
| Lakota | 374 (184, 761) | 1.7 (0.8, 3.5) | **2** |
| Man | 680 (503, 914) | 1.9 (1.4, 2.6) | **2.8** |
| Odienne | 261 (110, 630) | 2.8 (1.2, 6.7) | **1.7** |
| Oume | 540 (354, 829) | 1.9 (1.2, 2.9) | **2.5** |
| Sakassou | 262 (111, 634) | 2.7 (1.1, 6.6) | **1.7** |
| Sassandra | 1037 (647, 1684) | 3.2 (2, 5.2) | **3.5** |
| Seguela | 328 (141, 755) | 1.6 (0.7, 3.6) | **1.8** |
| Sinfra | 525 (343, 806) | 2.1 (1.4, 3.2) | **2.5** |
| Toumodi | 488 (313, 763) | 3.8 (2.4, 5.9) | **2.6** |
| Vavoua | 278 (80, 948) | 0.6 (0.2, 2.2) | **1.3** |
| Zuenoula | 214 (57, 783) | 1 (0.3, 3.5) | **1.1** |
| Bettie | 525 (179, 2025) | 8.9 (3, 34.4) | 3 (1.6, 4.3) |
| Blolequin | 597 (204, 1859) | 4.1 (1.4, 12.7) | 2.9 (1.6, 4.2) |
| Bocanda | 473 (153, 1342) | 3.8 (1.2, 10.8) | 2.6 (1.2, 3.9) |
| Botro | 612 (218, 2234) | 7.9 (2.8, 29) | 3.1 (1.8, 4.5) |
| Didievi | 434 (136, 1345) | 4.9 (1.5, 15.2) | 2.5 (1.2, 3.9) |
| Dikodougou | 415 (129, 1253) | 5.3 (1.6, 15.9) | 2.5 (1.2, 3.8) |
| Djekanou | 261 (81, 890) | 9.5 (3, 32.4) | 2.6 (1.3, 3.9) |
| Doropo | 328 (89, 945) | 5 (1.4, 14.4) | 2.2 (0.8, 3.5) |
| Fresco | 527 (180, 1695) | 4.9 (1.7, 15.8) | 2.8 (1.4, 4.1) |
| Gbeleban | 111 (28, 344) | 6 (1.5, 18.5) | 1.9 (0.5, 3.2) |
| Guitry | 474 (140, 1309) | 3 (0.9, 8.4) | 2.5 (1.2, 3.8) |
| Kani | 304 (79, 849) | 3.9 (1, 11) | 2 (0.7, 3.3) |
| Kouassi-kouassikro | 263 (82, 921) | 9.3 (2.9, 32.4) | 2.6 (1.2, 3.9) |
| M'bengue | 363 (103, 999) | 4.1 (1.2, 11.3) | 2.3 (1, 3.6) |
| Madinani | 255 (66, 776) | 6.4 (1.7, 19.4) | 2 (0.6, 3.3) |
| Nassian | 347 (102, 1150) | 7.6 (2.2, 25.3) | 2.4 (1.1, 3.7) |
| Niakaramandougou | 636 (229, 1990) | 4.6 (1.7, 14.4) | 3 (1.7, 4.3) |
| Samatiguila | 106 (25, 327) | 5.8 (1.4, 18.1) | 1.8 (0.5, 3.2) |
| Seguelon | 166 (43, 514) | 6.3 (1.7, 19.6) | 2 (0.6, 3.3) |
| Sikensi | 547 (195, 1838) | 6.7 (2.4, 22.5) | 2.9 (1.6, 4.3) |
| Sinematiali | 363 (110, 1114) | 6.4 (2, 19.7) | 2.4 (1.1, 3.7) |
| Sipilou | 285 (76, 873) | 6.3 (1.7, 19.2) | 2.1 (0.7, 3.4) |
| Taabo | 371 (116, 1122) | 6.2 (1.9, 18.8) | 2.4 (1.1, 3.7) |
| Tai | 567 (193, 1822) | 5 (1.7, 16.2) | 2.9 (1.5, 4.2) |
| Tehini | 310 (88, 996) | 7.5 (2.1, 24.1) | 2.2 (0.9, 3.6) |
| Tiassale | 550 (178, 1557) | 2.8 (0.9, 7.9) | 2.7 (1.3, 4) |
| Tiebissou | 443 (139, 1277) | 4.4 (1.4, 12.7) | 2.5 (1.2, 3.8) |
| Toulepleu | 459 (148, 1554) | 7.4 (2.4, 24.9) | 2.7 (1.4, 4.1) |
| Zouan-hounien | 511 (157, 1350) | 2.6 (0.8, 6.8) | 2.5 (1.2, 3.8) |
| Zoukougbeu | 342 (87, 905) | 2.8 (0.7, 7.3) | 2 (0.7, 3.4) |

**Figure 6:**
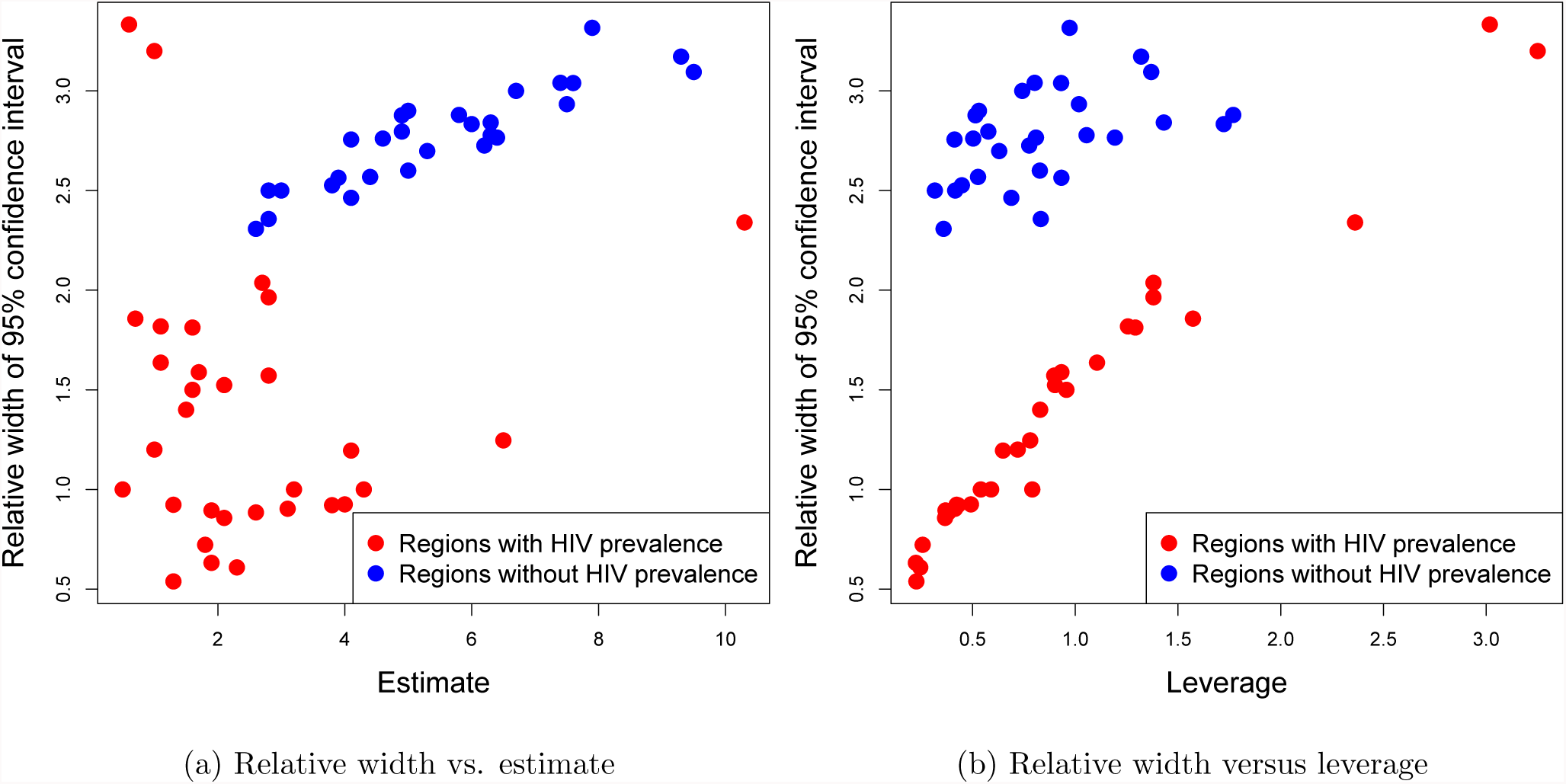
Impact of imputing HIV on uncertainty: Plots of (a) relative 95% CI widths versus estimates and (b) relative 95% CI widths versus the leverages leverages. Red and blue dots correspond to respectively regions with and without HIV prevalence.

In terms of absolute numbers, Abidjan has by far the highest predicted MSM population size although it has one of the lowest percentages. Both occurrences are due to the massive population of Abidjan. Outside of the five data regions, Katiola and Sassandra has the highest predicted MSM population size whereas some regions has predicted numbers as low as 100.

## 5 Discussion

We have provided fully Bayesian predictions combined with meaningful credible bounds for the population size of MSM in the 61 regions prioritized for HIV prevention and treatment services in Cˆote d’Ivoire. We have used the RDS based estimates as raw data and treated the associated confidence intervals as constant weights for the variances in regression. While it is common in survey sampling to treat an estimate of the sampling variance as known, in philosophy, this practice departs from proper modeling norms. More sophisticated models can be conceived where the data for each region is envisioned as a triplet consisting of the estimate, the upper and the lower quantiles. Directly using the individual survey data as an input to the model would be an even more fundamental approach. However, incorporating the RDS network into a hierarchical Bayesian area-level model remains a challenging problem.

The missing covariates adds to complications in statistical model evaluation of the two-step model. We evaluated the covariate selection and HIV prevalence imputation separately, before coalescing these two parts into the hierarchical model. Since, the spatial GP model induces dependence among all the observations, commonly used proper scoring rules (Gneiting & Raftery 2007) based on conditional independence, do not apply here. Further research needs to be conducted to come up with a proper scoring rule to evaluate the joint model in (2).

Bao et al. (2015) have demonstrated how to estimate populations sizes by incorporating data from multiple surveys and other data sources, in a fully Bayesian setup. While we have multiple estimates for each region, all of them are based on RDS and it is not clear how to adapt that approach when working with estimates from RDS. This once again highlights the need for more research on properly using RDS data in hierarchical models. Other relevant datasets, like MSM populations in other countries, if available, can be potentially leveraged to borrow strength in parameter estimation. However, care has to be taken when leveraging data from other countries, as different countries often have entirely different key population dynamics and borrowing strength may not be meaningful. Perhaps, more useful will be data for other associated key populations like Female Sex Workers, for the same set of regions. The correlation can be exploited in a multivariate setup to improve estimation of both populations.

The small sample size of numbers of cities with population size data for MSM has been a major limiting factor in modeling. However, the predictions presented here from this model represent the first empirically calculated estimates of the numbers of MSM in all areas of Cˆote d’Ivoire prioritized in the HIV response. This scenario of limited centers with measured population size is also not uncommon in the areas of the world where HIV prevalence is the highest given that these settings often also tend to criminalize same-sex practices or at least have significant stigma affecting MSM. In Southern and Eastern Africa, there is often only HIV prevalence data and size estimate data in one or a few urban centers for MSM though where studied, the HIV prevention and treatment needs are significant across these countries. While Cˆote d’Ivoire is in West Africa, it has one of the larger HIV epidemics in the region though limited information has been traditionally available for the numbers of MSM and the HIV burden among them. Thus, while the estimates provided here require further validation by supporting data to be collected in additional centers for MSM where predictions were completed, in the interim, these estimates can support the planning of the scale and content of HIV prevention and treatment programs for MSM in Cˆote d’Ivoire. Specifically, areas with wide credible intervals should be targeted for future surveys to improve modeling precision. Subsequently, validation and additional data points, will highlight the strengths and weaknesses of the current approach and pave the way for modeling improvements.

## Funding and acknowledgment

We would like to thank the participants of these studies and community groups that support them for engaging in the study. Data collection in Cˆote d’Ivoire was funded by the Global Fund to Fight AIDS, Tuberculosis and Malaria through the Government of Cˆote d’Ivoire National AIDS Control Program (PNPEC) contract to Enda Sant´e, an organization based in Senegal, and subcontracted for technical assistance to Johns Hopkins University. AD’s effort for methods development was supported by Project SOAR (Supporting Operational AIDS Research), Cooperative Agreement AID-OAA-A-14-00060, is made possible by the generous support of the American people through the President’s Emergency Plan for AIDS Relief (PEPFAR) and the United States Agency for International Development (USAID).

However, the contents of this article are the sole responsibility of Project SOAR, the Population Council, and the authors and do not necessarily reflect the views of PEPFAR, USAID, or the United States Government. SB’s effort was supported from the Johns Hopkins University Center for AIDS Research, an NIH funded program (P30AI094189), which is supported by the following NIH Cofunding and Participating Institutes and Centers: National Institute of Allergy and Infectious Diseases (NIAID), National Cancer Institute (NCI), National Institute of Child Health and Human Development (NICHD), National Heart, Lung, and Blood Institute (NHLBI), National Institute on Drug Abuse (NIDA), National Institute of Mental Health (NIMH), National Institute on Aging (NIA), Fogarty International Center (FIC), National Institute of General Medical Sciences (NIGMS), National Institute of Diabetes and Digestive and Kidney Diseases (NIDDK), and the Office of AIDS Research (OAR). The funders had no role in study design, data collection and analysis, decision to publish, or preparation of the manuscript.

## APPENDIX A RDS questions for multiplier methods

Questions asked of MSM recruited to participant in an RDS survey in order to provide an independent source of data for size estimation in Cˆote d’Ivoire in 2014

- **Unique object:** “Did you receive this object before?” [show single object]
- **NGO membership:** “Are you a member of the NGO Rainbow Plus, or have you ever participated one of their activities or even was hit by one of their peer educators?”
- “Are you a member of the NGO Alternative CI, or have you ever participated one of their activities or even was hit by one of their peer educators?”
- **Service:** “Have you received care at the Clinique de Confiance through the year 2014?”
- **Social event:** “Have you participated in the social event called “evening GNARA” which took place on Saturday, March 21, 2015 to space Embassy located in the Riviera II?”

